# CRISPR/Cas9-mediated gene deletion of the *ompA gene* in symbiotic *Enterobacter* impairs biofilm formation and reduces gut colonization of *Aedes aegypti* mosquitoes

**DOI:** 10.1101/389957

**Authors:** Shivanand Hegde, Pornjarim Nilyanimit, Elena Kozlova, Hema P. Narra, Sanjeev K. Sahni, Grant L. Hughes

## Abstract

**Background:** Symbiotic bacteria are pervasive in mosquitoes and their presence can influence many host phenotypes that affect vectoral capacity. While it is evident that environmental and host genetic factors contribute in shaping the microbiome of mosquitoes, we have a poor understanding regarding how bacterial genetics affects colonization of the mosquito gut. The CRISPR/Cas9 gene editing system is a powerful tool to alter bacterial genomes facilitating investigations into host-microbe interactions but has yet to be applied to insect symbionts.

**Methodology/Principal Findings:** To investigate the role of bacterial factors in mosquito biology and in colonization of mosquitoes we used CRISPR/Cas9 gene editing system to mutate the outer membrane protein A (*ompA*) gene of an *Enterobacter* symbiont isolated from *Aedes* mosquitoes. The *ompA* mutant had an impaired ability to form biofilms and poorly infected *Ae. aegypti* when reared in a mono-association under gnotobiotic conditions. In adults the mutant had a significantly reduced infection prevalence compared to the wild type or complement strains, while no differences in prevalence were seen in larvae, suggesting bacterial genetic factors are particularly important for adult gut colonization. We also used the CRISPR/Cas9 system to integrate genes (antibiotic resistance and fluorescent markers) into these symbionts genome and demonstrated that these genes were functional in vitro and in vivo.

**Conclusions/Significance:** Our results shed insights onto the role of *ompA* gene in host-microbe interactions in *Ae. aegypti* and confirm that CRISPR/Cas9 gene editing can be employed for genetic manipulation of non-model gut microbes. The ability to use this technology for site-specific integration of genes into the symbiont will facilitate the development of paratransgenic control strategies to interfere with arboviral pathogens such Chikungunya, dengue, Zika and Yellow fever viruses transmitted by *Aedes* mosquitoes.

**Author summary:** Microbiota profoundly affect their host but few studies have investigated the role of bacterial genetics in host-microbe interactions in mosquitoes. Here we applied the CRISPR/Cas9 gene editing system to knock out a membrane protein in *Enterobacter*, which is a dominant member of the mosquito microbiome. The mutant strain lacked the capacity to form biofilms, infected larvae and adults at lower titers, and had a reduced prevalence in adults. The lower prevalence in adults, but not larvae, likely reflects the difference in the modes of bacterial acquisition from the larval water of these two life stages. Importantly from an applied perspective, we also demonstrated that this editing technology can be harnessed for site-specific integration of genes into the bacterial chromosome. In proof-of-principle studies we integrated either a fluorescent protein or gene conferring antibiotic resistance into the bacterial genome and showed these transgenes were functional in mosquitoes. The specificity, flexibility, and simplicity of this editing approach in non-model bacteria will be useful for developing novel symbiotic control strategies to control arthropod-borne disease.

## Introduction

Mosquitoes harbor a community of microbes within their guts. In general, the gut-associated microbiome of mosquitoes tends to have low species richness but can differ greatly between individuals and habitats [1–8]. Importantly, these microbes can modulate many host phenotypes, several of which can influence vectorial capacity [9–11]. As such, it is imperative that we understand how the microbiome is acquired and maintained within mosquito vectors. While environmental factors unquestionably influences the mosquito microbiome composition and abundance [2–4, 8], studies are elucidating the role of microbial interactions[5, 7, 12, 13] and host genetic factors [14–18] in shaping the microbiome. However, we have a poor understanding of bacterial factors that influence colonization of the mosquito gut and this is likely an underappreciated force influencing host-microbe interactions in mosquitoes.

In other invertebrates, several bacterial genes have been implicated in gut colonization. For example, a genome wide screen exploiting transposon-sequencing found a suite of genes from the bacterium *Snodgrasselia* involved in colonization of the honey bee gut [19]. These bacterial genes were classified into the broad categories of extracellular interactions, metabolism and stress response [19]. Knock out of a purine biosynthesis gene in *Burkholderia* impaired biofilm formation and reduced bacterial colonization rates in a bean bug [20]. Biofilm formation was also shown to play a role in virulence of pathogenic *Pseudomonas* in artificial infections of *Drosophila*, with strains that lacked the capacity to form biofilms being more virulence to the host, while a hyperbiofilm strain was less virulent than the WT strain [21]. In other blood feeding invertebrates, bacterial genetics also appears critical for host colonization. Knockout of the type II secretion system in *Aeromonas veronii* reduced infection in *Hirudo verbena* leeches [22]. In Tsetse flies, the outer-membrane protein A (*ompA*) gene of *Sodalis glossinidius* is essential for symbiotic interactions [23]. *Sodalis* mutants lacking the *ompA* gene poorly colonized the fly gut compared to the wild type (WT) symbionts [23] and the mutant strain also had a reduced capacity to form biofilms [24]. Heterologous expression of the *ompA* gene from pathogenic *Escherichia coli* in *Sodalis* mutants induced mortality in the fly implicating this gene as a virulence factor in pathogenic bacteria [23]. Taken together, these studies suggest that bacterial genetic factors are critical for host colonization of invertebrates and that biofilm formation facilitates symbiotic associations in insects.

In mosquitoes, few studies have investigated how bacterial genetics affect gut colonization. However, evidence from experimental evolution studies suggests bacterial genetics plays a critical role. In two separate studies, *Enterobacter* was selected for increased persistence in the gut of *Anopheles gambiae* mosquitoes, the major malaria vector in sub-Saharan Africa, by repeatedly infecting mosquitoes with strains that persisted in the gut for longer periods of time [25, 26]. Transcriptomics comparisons of effective and ineffective colonizers in liquid media identified 41 genes that were differentially expressed between these two strains [26], further implicating the importance of bacterial genetics in mosquito infection, however the role of these genes in colonization of the mosquito gut has not been resolved. In a separate study, *in vitro* screening of a transposon mutant library of *Enterobacter* identified a *waaL* gene mutant that was insensitive to oxidative stress [27]. The *waaL* gene encodes an O antigen ligase which is needed for attachment of the O antigen to lipopolysaccharide and the mutant was found to have lower rates of colonization of the midguts of *Anopheles* mosquitoes [27].

Gene knockouts approaches in bacteria provide compelling evidence of the role of bacterial genes in host-microbe interactions [22–24, 27–29]. In general, most studies use transposon mutagenesis for gene knockout, which requires screening of the mutant library. A targeted gene knockout approach is highly desirable to investigate the functionality of bacterial genes in host-microbe interactions. In the past few years, the CRISPR/Cas9 gene editing system has been employed to modify bacterial genomes [30–32]. While much of the work has been done in model bacterial species [31–37], editing approaches have expanded into non-model bacterial systems [38–43]. Despite this expansion, the approach has been used less frequently used for host-associated microbes [39, 44], and rarely for arthropod symbionts. In the vector biology field, gene knockout approaches can be used to interrogate the role of bacterial genes responsible for host-microbe interactions, while the ability to integrate genes into the bacterial symbiont genome has great potential for applied paratransgenic control strategies [10, 45–47]. Previously, manipulation of non-model symbionts that associate with insect vectors have has been accomplished by plasmid transformation [48–55] or stable transformation of the genome using transposons or integrative plasmids [56–63], but the use of CRISPR/Cas9 gene editing in insect gut symbionts has yet to be accomplished. For paratransgenic strategies, stable site-specific integration of transgenes into the symbiont genome is critical, and as such, the application of CRISPR/Cas9 gene editing technology to non-model bacteria that associate with insect vectors will stimulate research in this field.

We therefore undertook studies to develop CRISPR/Cas9 genome editing approaches in an *Enterobacter* species isolated from *Aedes* mosquitoes. We used the <u>S</u>carless <u>C</u>as9 <u>A</u>ssisted <u>R</u>ecombineering (no-SCAR) method to disrupt the *ompA* gene of the non-model *Enterobacter* species [35]. After characterization of the mutant *in vitro*, we examined the role of the *ompA* gene in host-microbe interactions by re-infecting bacteria into mosquito in a mono-association. To demonstrate that the CRISPR/Cas9 gene-editing system could be useful for applied symbiotic control approaches we inserted genes conferring antibiotic resistance or a fluorescent protein into the bacterial genome and re-infected the altered strains back into mosquitoes. Our result sheds insights into the role of the *ompA* gene in host-microbe interactions in *Ae. aegypti* and confirm that CRISPR/Cas9 gene editing can be a powerful tool for genetic manipulation of native gut-associated microbes of mosquitoes.

## Results

### *Enterobacter* biofilm formation in *Ae. aegypti* guts

Over the course of conducting mono-axenic infections in *Ae. aegypti* mosquitoes with an *Enterobacter* symbiont, we repeatedly observed a conglomeration of bacterial cells in the gut that was indicative of a biofilm (Figure 1, Figure S1 A-C). This formation of bacteria has a similar appearance to biofilms observed in the guts of other insects [21, 24]. No bacteria were observed in the intestinal track of *Ae. aegypti* when infected with *E. coli* (Figure 1, S1 D-F), although as seen previously, infection with this bacterium enabled mosquito development [64]. We therefore examined the role of bacterial genetics in biofilm formation and host colonization of this gut-associated bacterium of *Aedes* mosquitoes. While several genes have been implicated in biofilm formation [21, 24], we chose to knockout the *ompA* gene of *Enterobacter* given that this gene has been demonstrated to influence biofilm formation and gut colonization of *Sodalis* [23, 24], an *Enterobacteriaceae* symbiont of Tsetse flies. We used the CRISRP/Cas9 genome editing system to mutate the symbionts genome.

**Figure 1.**
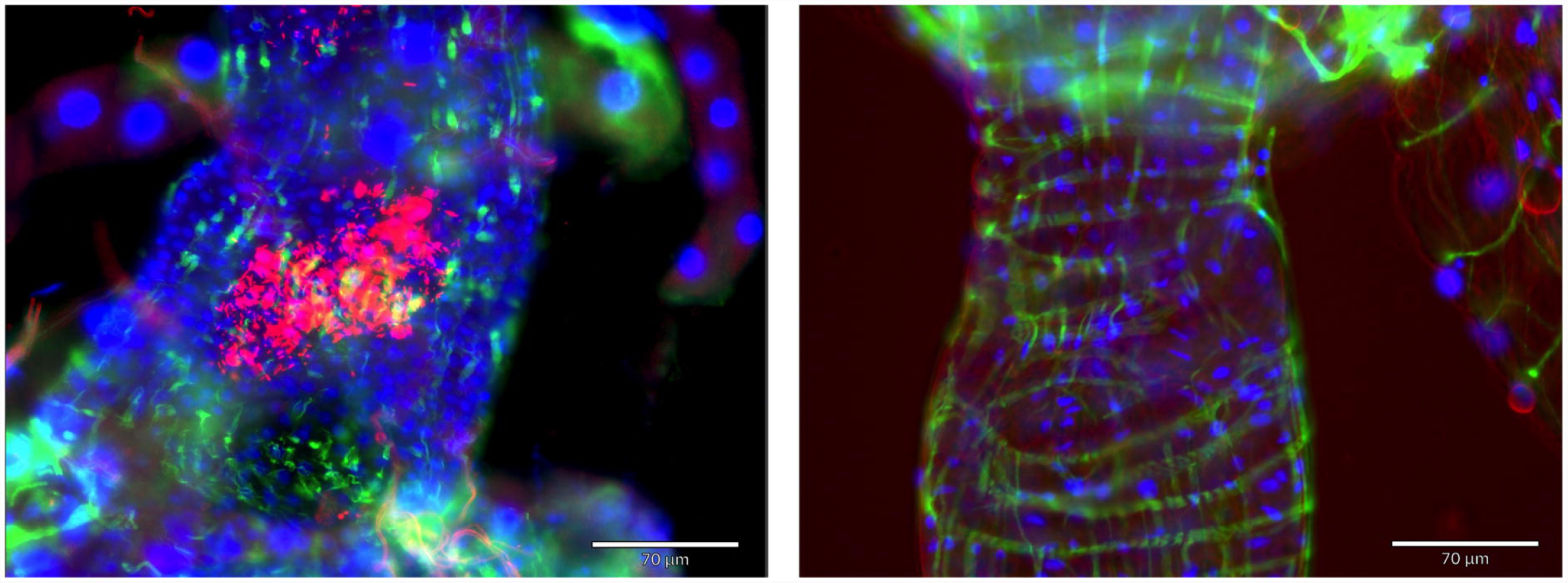
Midgut infection of *Enterobacter* and *E. coli* in mono-associations of *Aedes* mosquitoes. *Enterobacter* forms a biofilm in the gut of *Aedes aegypti* mosquitoes (left) while no bacteria were observed in the gut of mosquitoes reared with *E. coli* under gnotobiotic conditions (right). Bacteria possessed a plasmid expressing mCherry. Blue – host nuclei. Green – host actin cytoskeleton stained with phalloidin. The scale bar is 70 µm.

### Genome editing in *Enterobacter* bacteria isolated from mosquitoes

To edit the *Enterobacter* isolate that resides within the gut of *Aedes* mosquitoes, we employed the no-SCAR gene editing approach that had been developed in *E. coli* [35]. To optimize the approach in our hands, we performed initial experiments in *E. coli* to delete a ∼ 1 kb region of the *ompA* gene (Figure 2A). As the no-SCAR approach exploits the λ-Red recombineering system to repair double stranded breaks, we transformed bacteria with a double stranded DNA template that had regions of homology flanking the gRNA site (250 bp for each arm). Using this approach, we successfully deleted a 1001 bp fragment of the *ompA* gene. From the colonies we screened, we saw an editing at a frequency of 6.25% (N = 48) (Figure 2A). For *Enterobacter*, we altered our editing procedure to delete a 598 bp fragment from the *ompA* gene. This was done to enhance the efficiency of obtaining mutants [35] and accommodate the PAM site which was at a different location in the *ompA* gene in *Enterobacter*. Using a donor template designed for the *Enterobacter ompA* gene that had similar length flanking homology arms as the previous experiment done in *E. coli*, we obtained mutant knockouts at a rate of 32% (N = 50) (Figure 2B). For both bacterial species, Sanger sequencing across the integration site indicated the deletion occurred at the expected loci in the bacterial genome (Figure 2C; S1 Appendix).

**Figure 2.**
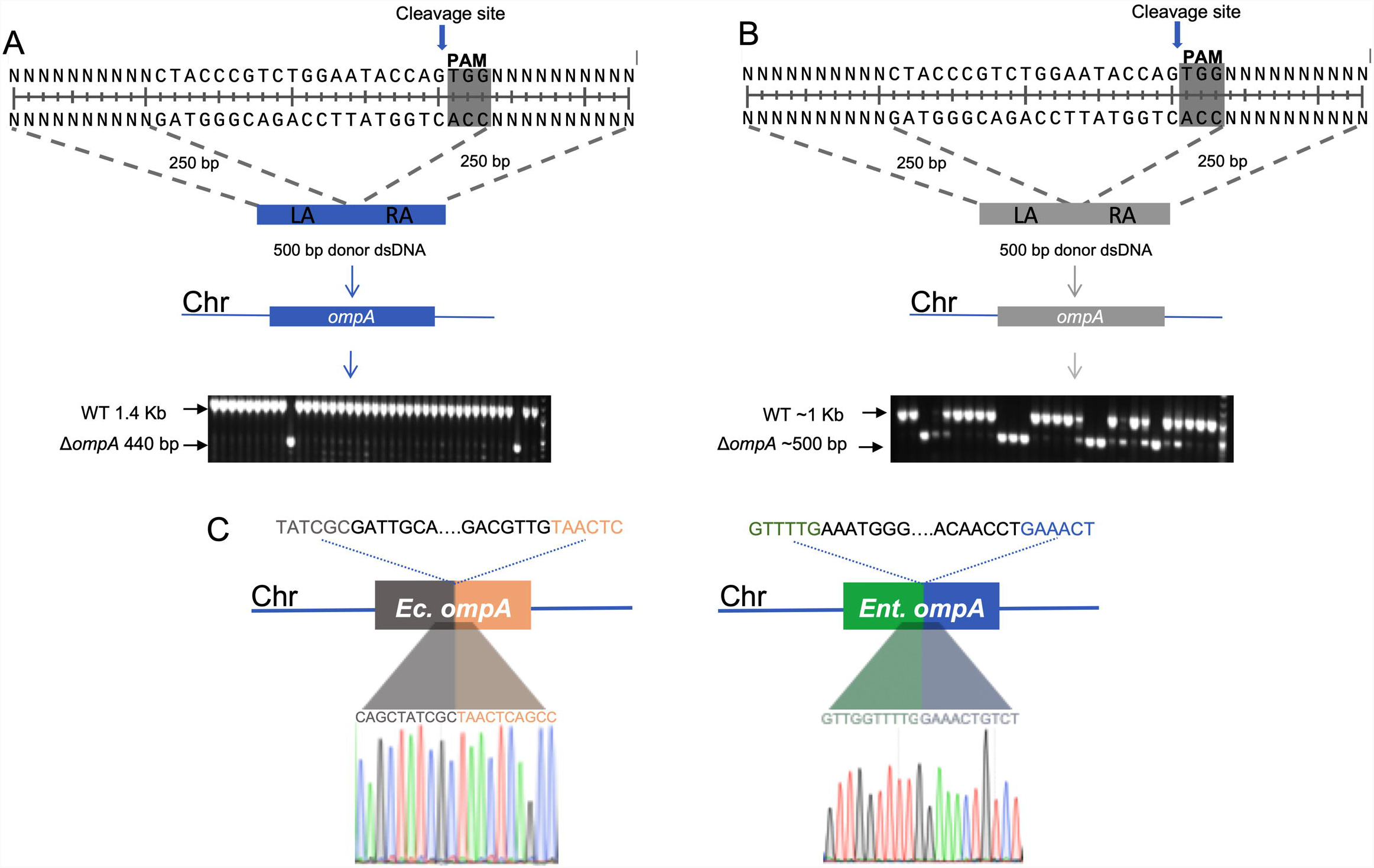
CRISPR/Cas9 genome editing in bacteria. A schematic of the editing approach and screening of putative mutants in *E. coli* (A) and *Enterobacter* (B). A ∼ 1kb fragment of *E. coli* BL21(DE3) was deleted using no-SCAR protocol. The 250 bp of left arm (LA) and right arm (RA) was assembled to generate 500 bp donor DNA. The transformants were screened via colony PCR with primers binding in regions flanking the deletion. Similar to strategy employed in *E. coli*, knockout of the *ompA* gene from *Enterobacter* isolated from the mosquito gut was created by deleting the 598 bp fragment. The grayed area indicates the PAM site in the *ompA* gene and arrow shows cleavage site in the genome. (C) The sequence of the *ompA* mutation in *E. coli* and *Enterobacter* was confirmed by Sanger sequencing. The sequence above the gene within the dotted line has been deleted. The chromatogram shows the 10 bp flanking the deletion.

### Characterization of the *Enterobacter ompA* mutant

We quantified the growth rates of the Δ*ompA* mutant in comparison to the WT *Enterobacter* and the Δ*ompA/ompA* complement in liquid LB media. We saw minimal differences in the growth between the WT, the Δ*ompA* mutant or the Δ*ompA/ompA* complement (Figure 3A). To examine the stability of the deletion, we subcultured the Δ*ompA* mutant on LB media for 10 generations and performed PCR to amplify across the deletion. At alternative generations PCR analysis indicated the deletion was present indicating genomic stability at this site (Figure 3B).

**Figure 3.**
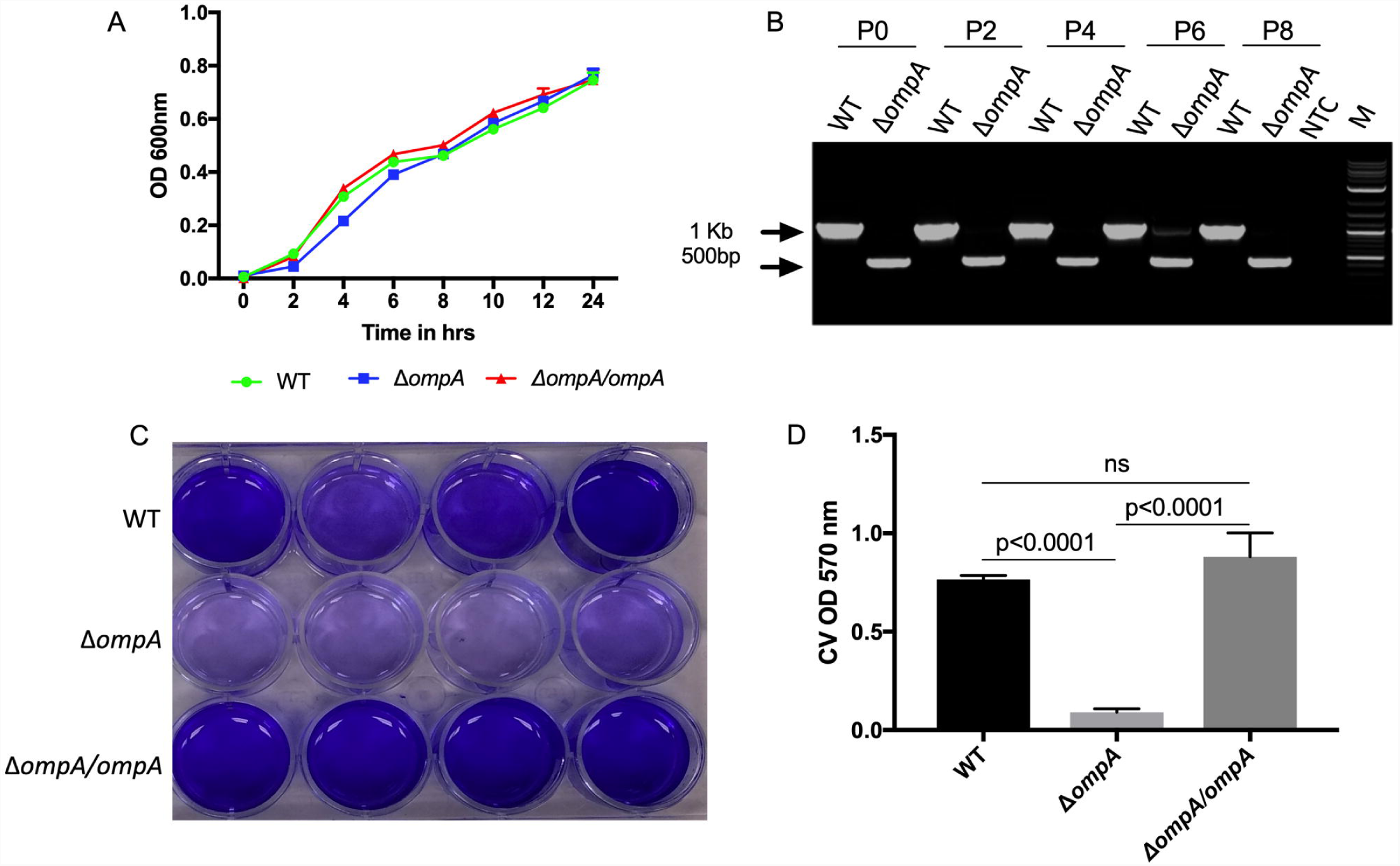
In vitro characterization of the *ompA* mutation. The *Enterobacter* Δ*ompA* mutant had a similar growth rate compared to both the WT and the Δ*ompA/ompA* complement in liquid LB media (A). The stability of mutant was evaluated *in vitro* by continuous subculturing in LB medium (B). Genomic DNA of alternative subcultures was used as template for PCR using gene specific primers that amplified across the deletion. Two separate gel images were merged to create the figure 2B. Passage 8 was run on a separate gel to passages 0 – 6. Biofilm formation was assessed using the CV biofilm assay for the WT, Δ*ompA* mutant and the Δ*ompA/ompA* complement (C). Quantification of the relative biofilm formation normalized by the number of bacteria per well (D).

Previously, *ompA* has been shown to be important in biofilm formation as *Sodalis* deletion mutants were unable to form biofilms [24]. As such we characterized *in vitro* biofilm formation using the crystal violet (CV) biofilm assay. After visual inspection, it was clear the Δ*ompA* mutant had distinctly less biofilm deposition compared to either the WT or the Δ*ompA/ompA* complement (Figure 3C), and after quantification and normalization to account for any difference in growth between the strains, biofilm formation was confirmed to be significantly different between the Δ*ompA* mutant and the WT (Figure 3D; Tukey’s multiple comparisons test, P < 0.0001) or Δ*ompA/ompA* complement (Tukey’s multiple comparisons test, P < 0.0001), while there was no significant differences between the WT and the Δ*ompA/ompA* complement (Tukey’s multiple comparisons test P = 0.2).

### The role of *ompA* gene in mosquito infection

To examine the importance of the *ompA* gene on bacterial colonization of mosquitoes, we infected *Ae. aegypti* mosquitoes in a mono-association under gnotobiotic conditions [64]. This infection method was used to avoid other gut-associated microbes influencing host colonization rates [7] and it also enable straightforward quantification of introduced bacteria by measuring colony forming units (CFUs). In larvae we saw a significant reduction in bacterial titer in the mutant compared to both the WT (Kruskal-Wallis test; P < 0.01) and the Δ*ompA/ompA* complement (Kruskal-Wallis test; P < 0.05) (Figure 4A). Similarly, in adults, there was a significant reduction in bacterial infection in the Δ*ompA* mutant compare to either the WT or Δ*ompA/ompA* complement (Kruskal-Wallis test; P < 0.001) (Figure 4B). While no significant changes were seen in the prevalence of infection (number of mosquitoes infected) in the larval stage (Figure 4C, Fisher’s exact test; WT compared to Δ*ompA* P = 0.24 and Δ*ompA* compared to Δ*ompA/ompA* P = 0.24), in adults, the prevalence of infection was significantly different (Figure 4D, Fisher’s exact test; WT compared to Δ*ompA* P < 0.0001 and Δ*ompA* compared to Δ*ompA/ompA* P < 0.0001), with only 45% of adults infected by the Δ*ompA* mutant compared to 95% and 88% by the WT and Δ*ompA/ompA* complement, respectively. We also examined the growth rates of mosquitoes administered with the WT, Δ*ompA* mutant and Δ*ompA/ompA* complement. No significant differences were seen in the time to pupation (Figure 5A) or percentage of first instar larvae that reached adulthood (Figure 5B) between any of the strains.

**Figure 4.**
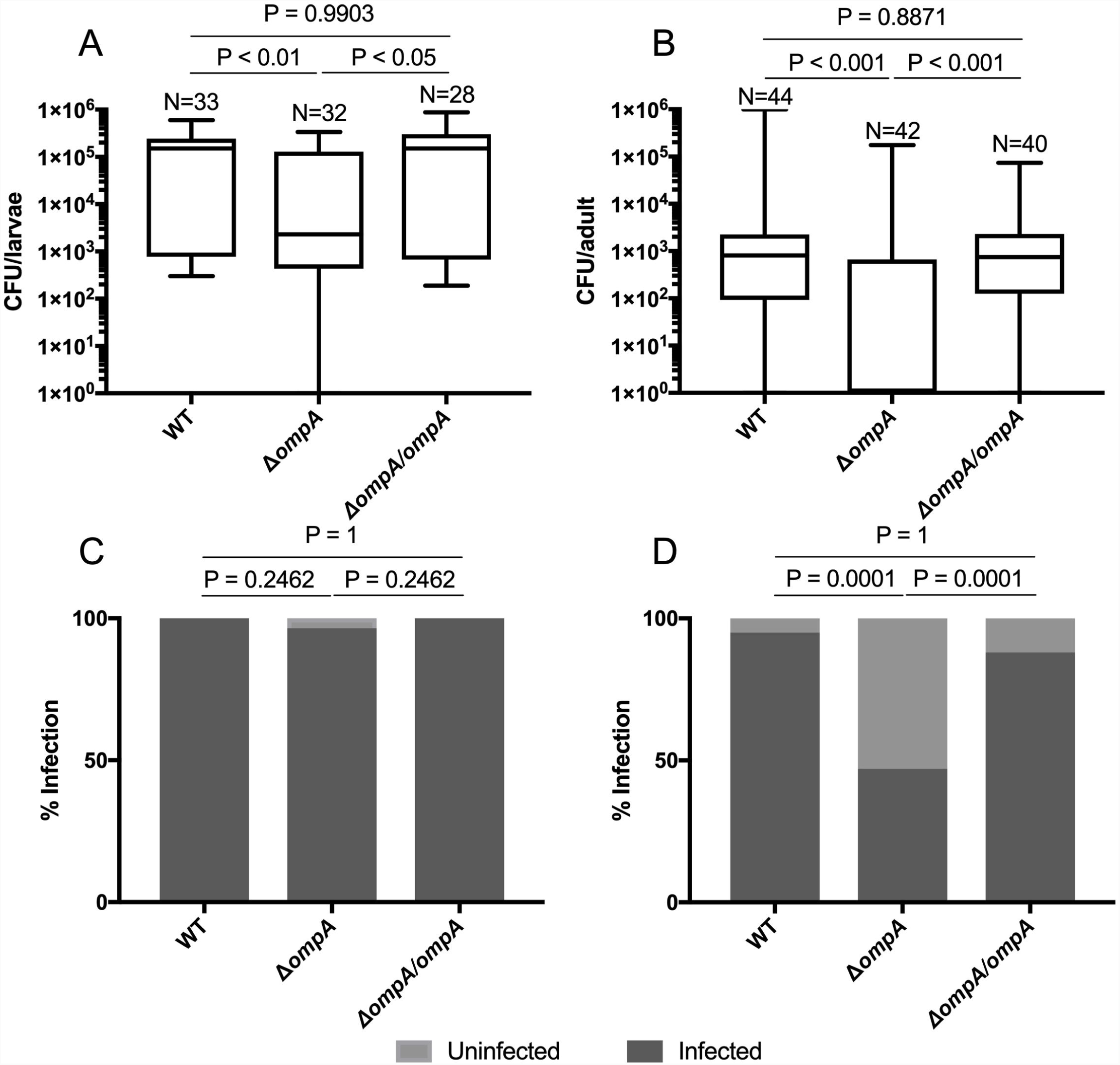
The Δ*ompA* mutant poorly infected mosquitoes. Infection of *Enterobacter* strains (WT, Δ*ompA* mutant and Δ*ompA/ompA* complement) reared in a mono-association using a gnotobiotic rearing approach for larvae (A) and adults (B). L4 and 3-4 days post emergence adults were screened for bacterial load by plating on LB media to quantify the bacteria. The prevalence of infection (number of mosquitoes infected) between the treatments was calculated comparing number of infected to uninfected larvae (C) or adults (D).

**Figure 5.**
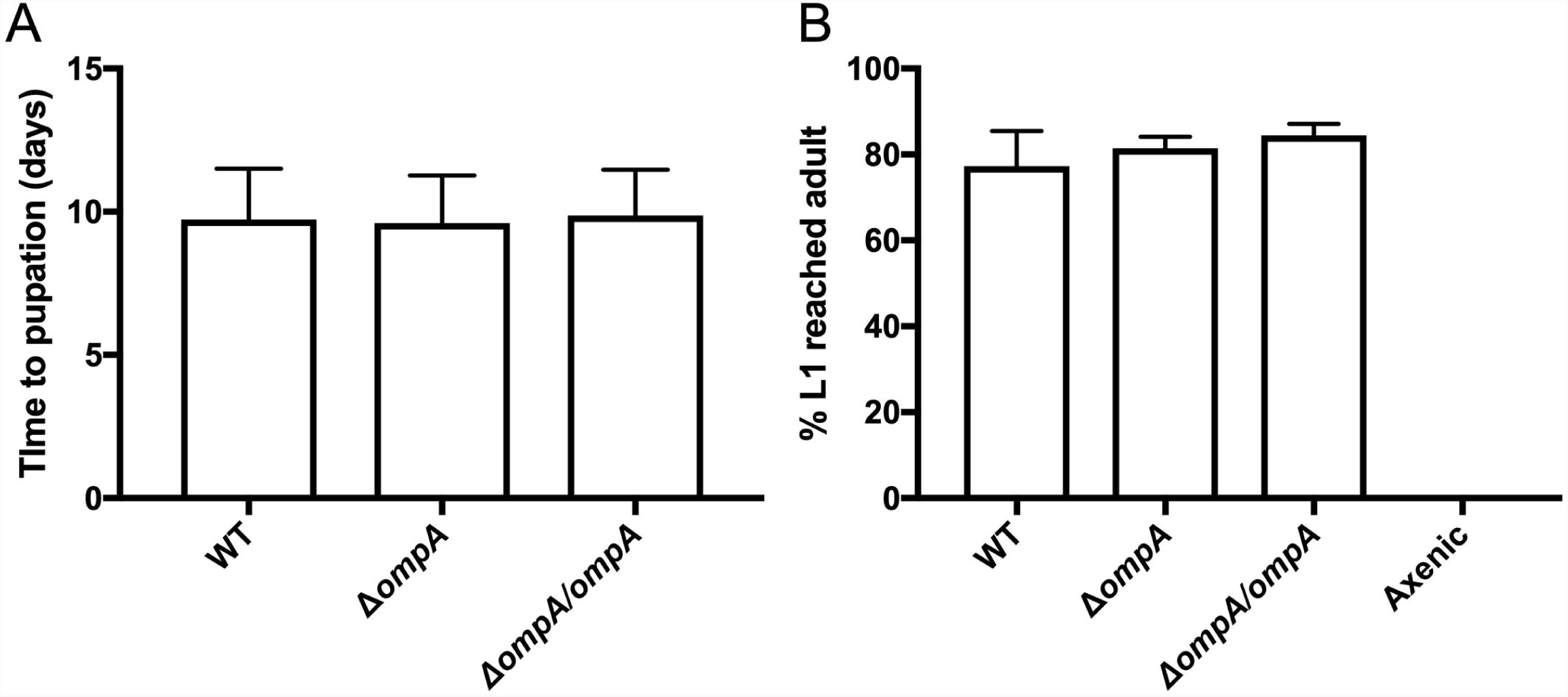
The Δ*ompA* mutant does not affect growth rates or development of mosquitoes. The growth rate (time to pupation) (A) and development (percentage of L1 larvae to reach adulthood) (B) was observed in mosquitoes infected with *Enterobacter* strains (WT, Δ*ompA* mutant and Δ*ompA/ompA* complement) reared in a mono-association.

### Integration of genes into the *Enterobacter* chromosome

We undertook experiments to demonstrate the CRISPR/Cas9 gene-editing approaches can be used to integrate genes into the chromosome of non-model bacteria that associate with mosquitoes. We created two independent transgenic strains that had either, a gene encoding mCherry fluorescence or a gene encoding resistance to the antibiotic gentamicin, inserted into the bacterial chromosome. These genes were integrated into the genome using the same gRNA that was used for deletional mutagenesis (Table S1), and as such, these insertions also disrupted the *ompA* gene. Sequencing across the integration site indicated the insertion of these genes occurred within the *ompA* gene and thereby disrupted its function (Figure 6A and 6D). Continual subculturing was undertaken for both strains and molecular analysis indicated the stability of these lines for ten generations (Figure 6B and 6E). To demonstrate the integrated genes were functional, we observed expression of mCherry fluorescence and successfully cultured the strain containing gentamicin resistance on plates containing the antibiotic (Figure 6C and 6F). Finally, we infected these transgenic strains into mosquitoes to demonstrate that these strains were able to colonize the mosquito gut and functionality of the integrated gene was confirmed by observing fluorescence or by rearing the *Enterobacter ompA*::gentamicin strain in mosquitoes administered sugar supplemented with gentamicin. Fluorescent bacteria were observed in the gut of mosquitoes while no signal was seen in controls (WT *Enterobacter* infected mosquitoes) (Figure 5G). The *Enterobacter ompA*::gentamicin was successfully rescued from mosquitoes reared on gentamicin and was seen to stably infect mosquitoes over time at a density of 1×10^4^ CFUs/mosquito. Consistent with our previous finding (Figure 4B), the WT bacteria initially infected mosquitoes at higher titers (T test; day 0 P < 0.001). However, at 4 days post infection (dpi), the total bacterial load of culturable microbes in mosquitoes supplemented with WT *Enterobacter* was significantly reduced when reared on sugar supplemented with antibiotic (T test; day 4 P < 0.05), and no CFUs were recovered after at 6 dpi (T test; day 6 P < 0.001) (Figure 6H).

**Figure 6.**
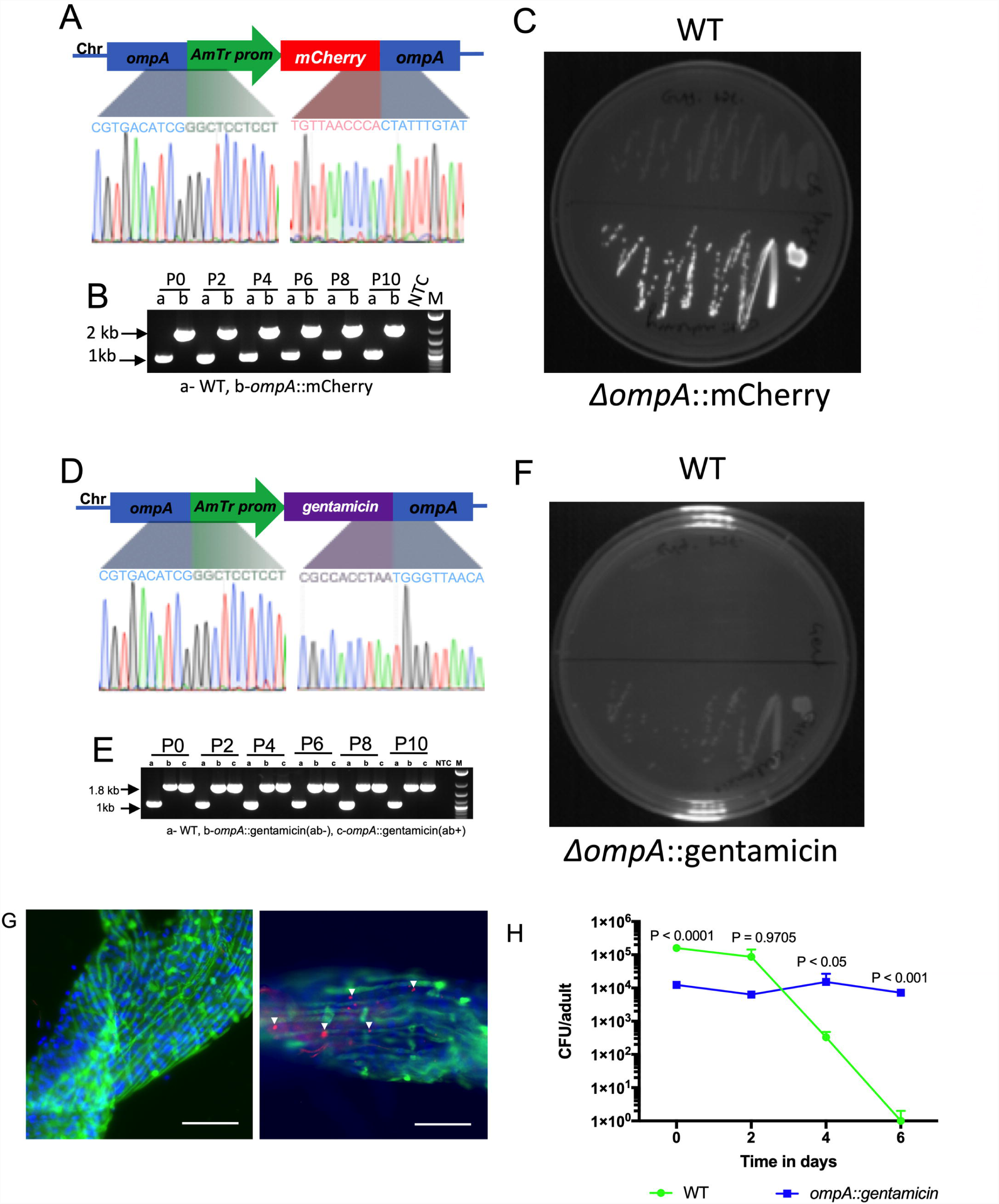
Integration of mCherry and gentamicin into the *Enterobacter* genome. Sanger sequence across the integration site, stability of the inserted gene and *in vitro* expression of the inserted gene for the Δ*ompA*::mCherry (A-C) and the Δ*ompA*:: gentamicin (D-F) strains. The chromatogram shows the sequence spanning the inserted sites. Strains were continually subcultured for 10 passages and PCR was done to examine the stability of the insert (B; Δ*ompA*::mCherry plus WT; E Δ*ompA*::gentamicin passaged with (ab+) or without (ab-) gentamicin plus WT). mCherry fluorescence or ability to grow on selective media containing gentamicin confirmed the expression of the transgene *in vitro*. Mosquitoes were inoculated with the *Enterobacter* strains to confirm expression of the transgene *in vivo*. Dissected midgut infected with Δ*ompA*::mCherry (left) or negative control (right; WT bacteria without expression plasmid) (G). Midguts were stained with phalloidin (green) and DAPI (blue). The scale bar is 30 µM. The WT and Δ*ompA*::gentamicin *Enterobacter* strains were fed to adult mosquitoes for 3 days in a sugar meal before gentamicin was administered to mosquitoes in a sugar meal (H). Mosquitoes were collected every second day and CFUs assessed. Pairwise comparisons were conducted at each time point using a T test (* −P < 0.05, *** P < 0.001, **** P < 0.0001).

## Discussion

We harnessed the CRISPR/Cas9 gene editing system to create knockout mutants in an *Enterobacter* gut symbiont of *Ae. aegypti* mosquitoes enabling us to examine the role of bacterial genetics, specifically the *ompA* gene, in biofilm formation and gut colonization. A deletion of the *ompA* gene of *Enterobacter* decreased bacterial colonization of the mosquito host at both the larval and adult stages after infection in a mono-association. Strikingly, we found this effect was most pronounced in adult mosquitoes with more than half of the mosquitoes not possessing any culturable mutants, while there was no difference in prevalence of infection between the mutant and WT bacteria in larvae. The reduced prevalence of mutant bacteria in adults likely reflects differences in microbial colonization of each mosquito life stage. Larvae are continually subjected to bacteria in the larval water habitat while adults only have a short time frame to acquire bacteria from the aquatic environment immediately after eclosion, when they are thought to imbibe a small amount of larval water which seeds the gut with microbiota [65]. Our data shows greater variation in colonization of the adult stage between the mutant and WT strains, indicating that the *ompA* gene, and potentially bacterial factors in general, may be critical for colonization of the adult gut. These findings are also consistent with other sequence-based studies, that indicate adult stages have greater variability in species composition of their microbiota, while the microbiome of immature stages is similar to the microbiota in larval water habitat [2–5, 8, 66].

Overall, our findings are similar to studies done in Tsetse flies whereby an *ompA* mutant of *Sodalis*, an *Enterobacteriaceae* symbiont, has impaired biofilm formation and reduced colonization rates [23, 24]. These studies, in conjunction with our work, suggests that the *ompA* gene is imperative for symbiotic associations within dipterans. It also suggests that biofilm formation may be a strategy employed by bacteria to colonize the gut of insects. In pathogenic infections in mammals, biofilms enable bacteria to colonize new niches, promote infection and are associated with virulence [67]. Although less is known regarding the importance of biofilm formation in insects, in an artificial *Pseudomonas-Drosophila* infection model, biofilm formation was associated with virulence and host survival [21]. In a natural symbiotic association between bean bugs and *Burkholderia*, disruption of a purine biosynthesis gene in the bacterium also reduce biofilm formation and colonization of the insect [20] In mosquitoes, gut biofilm formation could also have implications for vector competence as *Chromobacterium*, which was isolated from *Aedes* mosquitoes, produced molecules that inhibited dengue virus only when grown *in vitro* as a biofilm but not when grown in a planktonic state [68]. Despite this, it was unknown if biofilm formation occurred *in vivo* in the mosquito [68]. Our data provide evidence that biofilms occur within the gut of mosquitoes and facilitate host colonization.

While we have shown that the *ompA* gene of *Enterobacter* is important for host colonization, we see no evidence that deletion of this gene alters mosquito development or growth rates. This is in contrast to the *Riptortus*-*Burkholderia* symbiosis whereby mutation of the *purT* gene in *Burkholderia* resulted in reduced growth rates and reduction in body weight of the host compared to insects that were infected with the WT bacterium [20]. The difference in our study to the findings in the *Riptortus*-*Burkholderia* symbiosis could be related to different requirements of the bean bug compared to the mosquito host as well as the different genes mutated in the symbionts. Our findings are consistent with a previous study in *Ae. aegypti* whereby an *ompA* mutant of *E. coli* did not influence growth reared in a mono-association [69]. Using a similar gnotobiotic system that exploits the ability to sterilize mosquito eggs and rescue development by nutritional supplementation, several recent reports describe approaches to create bacteria-free mosquitoes [69, 70]. Here, we reared mosquitoes in a mono-association where they were only subjected to *Enterobacter.* However, more than half the adult mosquitoes inoculated with the Δ*ompA* mutant were not infected by bacteria, as evidenced by the inability to culture bacteria from these insects. Nevertheless, these mosquitoes had similar development and growth rates compared to mosquito possessing WT bacteria. The use of mutant bacteria that rescue development but have an impaired ability to colonize mosquitoes may provide a simple means to create axenic adult mosquitoes.

CRISPR/Cas9 gene editing has revolutionized genetic approaches in model and non-model bacteria [31–43]. However, there has been limited use of this technology in symbiotic microbes of arthropods. Here we demonstrate that editing approaches functional in *E. coli* can be easily applied with minimal adaptation to phylogenetically related symbiotic bacteria that associate within the guts of mosquitoes. The application of CRISPR/Cas9 genome editing to gut-associated bacteria of mosquitoes has significant applied potential. Paratransgenesis strategies are being evaluated in a range of medical and agricultural systems to mitigate pathogen transmission from insect vectors, however, most approaches engineer symbionts by plasmid transformation [49–55, 71] and where genome integration has been accomplished in symbionts [58–61], it has often been done with technologies that did not allow for site specific integration. Paratransgenic approaches suitable for use in the field will need to stably integrate genes into the bacterial genome in a manner that does not compromise bacterial fitness. Exploiting the flexibility and specificity of the CRISPR/Cas9 to integrate genes in intergenic regions of the bacterial chromosome will undoubtedly be beneficial for these applied approaches.

In summary, we have demonstrated that the CRISPR/Cas9 gene editing system can be applied to symbiotic bacteria that associate with eukaryotic hosts to interrogate the role of bacterial genes in host-microbe associations. We created knockout and knockin mutants by deleting and disrupting the *ompA* gene of *Enterobacter*. The knockout mutant displayed a reduced ability to form biofilms and colonize the gut of *Ae. aegypti* mosquitoes in a mono-association, demonstrating bacterial genetic factors are important determinants that influence colonization of mosquito guts. *Aedes* mosquitoes are becoming powerful systems to investigate the genetics of host-microbe interactions given the scientific community has simple and efficient approaches to alter both the microbes (this study) and mosquito host genome [72, 73] at their disposal, as well as methods to create mono-associated mosquito lines[7, 64]. Finally, rapid, efficient, and site specific gene editing approaches for gut bacteria that associate with mosquitoes will facilitate the development of novel paratransgenic approaches to control arthropod-borne disease [57].

## Experimental procedures

### Bacterial and mosquito strains

*E. coli* BL21(DE3) (NEB) and an *Enterobacter* strain previous isolated from a lab-reared colony of *Ae. albopictus* (Galveston) mosquitoes [7] were used in this study. Cultures were grown in liquid LB media at 37°C with the appropriate antibiotic unless stated otherwise. Mosquitoes were reared in the UTMB insectary under conventional conditions or in mono-associations (described below).

### CRISPR gene editing

Editing the *ompA* gene of *E. coli* and *Enterobacter* was complete as described in Reisch and Prather [35]. The protospacer sequence for the *ompA* gene was designed using the CHOPCHOP [74, 75], and cloned into pKDsgRNA-ack plasmid [35] directly upstream of gRNA scaffold using REPLACR mutagenesis protocol [76]. Two protospacer sequences were designed for each gene and the one which had lower escape rate after plating with or without aTC (S1 Table). The plasmids were acquired from Addgene (S2 Table; Addgene plasmid 62655 and 62654). The resulting plasmids pKDsgRNA-Ec-*ompA* and pKDsgRNA-Ent-*ompA* were Sanger sequenced to confirm insertion of protospacer sequence. These plasmids were then transformed into either *E. coli* or *Enterobacter* containing the pCas9-CR4 plasmid. Transformants were selected at 30°C on LB agar plate containing spectinomycin (50 µg/mL), chloramphenicol (34 µg/mL), and with or without anhydrotetracycline (aTC) at 100ng/mL. Colonies from the –aTC plate were grown overnight in LB broth with the appropriate antibiotic at 30°C. A 1:100 diluted overnight culture was (grown until 0.4 OD_600_) supplemented with 1.2% arabinose to induce the expression of λ-Red recombinase. Cells were then transformed with 1-1.5 µg of double stranded donor DNA that flanked the PAM site for homologous recombination. Donor DNA was created by either PCR amplification or by gene synthesis (Genewiz). Regardless of the method of construction, each donor had flanking regions of 250 bp homologous to the target DNA. The resulting colonies were screened for mutations by colony PCR with primers flanking the integration site and positive clones were Sanger sequenced (S3 Table). Positive colonies were grown in LB broth and genomic DNA was isolated. For further validation, the flanking regions of deletion or insertions were amplified and the PCR product Sanger sequenced.

### Stability of insertion

The stability of the knockout Δ*ompA* mutant and the knockin *ompA*::gentamicin and *ompA*::mCherry strains was assessed in LB medium. The *omp*A::mCherry and knockout Δ*ompA* mutant cultures were grown for 10 passages in LB broth. At each passage 40 µl of culture was transferred into 4ml fresh LB medium. The *ompA*::gentamicin strain was grown with or without gentamicin (50 µg/mL). Genomic DNA was isolated from the 0, 2, 4, 6, 8 and 10^th^ subculture and PCR that amplified across the integration site was performed.

### Complementation of *ompA* mutant

Functional rescue of the *omp*A mutation was achieved by complementing the mutant with the WT gene. The WT *omp*A gene was amplified from *Enterobacter* genomic DNA and cloned into the pRAM-mCherry vector^7^ and thereby creating pRAM-mCherry-*Ent*-*OmpA.* The Sanger sequence-verified plasmid was transformed into the Δ*ompA* mutant, thereby generating the Δ*ompA/ompA* complement strain. Colonies that acquired the plasmid were selected on LB plates containing kanamycin (50 µg/mL).

### *In vitro* characterization of *Enterobacter* strains

To assess the impact of the gene deletion on bacterial growth the WT, Δ*ompA* mutant and Δ*ompA/ompA* complement were grown in LB broth and the density of bacteria (OD_600_) was quantified by spectrophotometer. A 1:100 dilution of an overnight culture was inoculated into a 5 ml LB broth in 50 ml tube and incubated at 37°C for 24 hrs. At 2, 4, 6, 8, 10, 12 and 24 hours growth was recorded at OD_600._ The biofilm assay was performed as described previously [77, 78]. Briefly, biofilm formation by *Enterobacter* strains was quantified on polystyrene microtiter plates after 72 h of incubation at 37°C by CV staining. Three independent experiments were performed, and the data were represented as CV OD_570_ after normalizing by CFUs.

### Mosquito infections

Mono-association in *Ae. aegypti* mosquitoes were done using gnotobiotic infection procedure [7, 64],+ with slight modifications. Briefly, mosquito eggs were sterilized for 5 min in 70% ethanol, 3 min 3% bleach+0.01% Coverage Plus NPD (Steris Corp.), 5 min in 70% ethanol then rinsed three times in sterile water. Eggs were vacuumed hatched for 30-45 min and left overnight at room temperature to hatch any remaining eggs. Exactly twenty L1 larvae were transferred to T175 flask containing 60 ml of sterile water and fed on alternative days with 60 µl of fish food (1 µg/µl). Larvae were inoculated with 1×10^7^/ml of either the WT *Enterobacter*, the Δ*ompA* mutant or the Δ*ompA/ompA* complement. The WT and Δ*ompA* strains were transformed with the pRAM-mCherry plasmid that conferred resistance to kanamycin (but did not possess a functional *ompA* gene). L4 larvae were collected, washed three times with 1X PBS, and then homogenized in 500 µl of 1X PBS and 50 µl of homogenate was plated on LB agar containing 50 µg/mL kanamycin. Similarly, adult mosquitoes were collected 3-4 days post emergence and bacterial infection was quantified in the same manner as larvae. In order to assess the growth of the mosquitoes, time to pupation and growth rate were observed. Time to pupation was determined by quantifying the number of pupae each day post hatching, while survival to adulthood was calculated by quantifying the number of L1 larvae that reached adulthood. The experiment was repeated three times.

Knock-in mutants were administered to adult *Ae. aegypti* in a sugar meal. Three to four day old mosquitoes were fed with 1×10^7^ of WT or the Δ*ompA*::gentamicin strain for three days in 10% sucrose solution. After three days, mosquitoes were either administered sugar supplemented with gentamicin (50 µg/mL) or sugar without antibiotic. CFUs were determined at days 0, 2, 4, and 6 dpi by plating homogenized mosquitoes (N=10) on LB agar. Similarly, the Δ*ompA*::mCherry and WT *Enterobacter* were fed to mosquitoes and midguts were dissected to assess the colonization of bacteria in the tissue. For visualization of bacteria, midguts were fixed in 1% paraformaldehyde (PFA) in 1X PBS for 30 minutes and permeabilized with 0.01% Triton X-100 in 1X PBS for 20 min. The tissues were stained with 1:250 diluted Phalloidin (Sigma) for 20 minutes and samples were washed twice with 1X PBS for 10 minutes. Finally, midguts were then stained with 1:500 diluted DAPI (Invitrogen) for 10 min. Samples were transferred to slides and mounted with ProLong™ Gold Antifade (Invitrogen). The slides were observed under Revolve FL (ECHOLAB).

## Acknowledgements

We would like to thank the UTMB insectary core for providing mosquitoes. GLH was supported by NIH grants (R21AI138074, R21AI124452 and R21AI129507), a University of Texas Rising Star award, the Western Gulf Center of Excellence for Vector-borne Diseases (CDC grant CK17-005), the Robert J. and Helen Kleberg Foundation and the Gulf Coast Consortia. This work was also supported by a James W. McLaughlin postdoctoral fellowship at the University of Texas Medical Branch to SH and a Dutsadi Piphat Scholarship and Thailand Research Chair Grant from NSTDA (p-15-50004) to PN. We also wish to thank Dr Ulrike Munderloh (University of Minnesota) for the kind gift of the pRAM plasmid.

## Figure legends

**S1. Figure. Midgut infection of *Enterobacter* and *E. coli* in mono-associations of *Aedes* mosquitoes.** Different gut tissue locations showing the conglomeration of bacterial cells when infected in mono-association in *Aedes* mosquitoes with *Enterobacter* (A-C). However, infection of *E. coli* did not show any infection in the gut of the mosquitoes (D-F). The pictures were taken from different field of view of the gut dissected from different mosquitoes.

**S1. Table.**
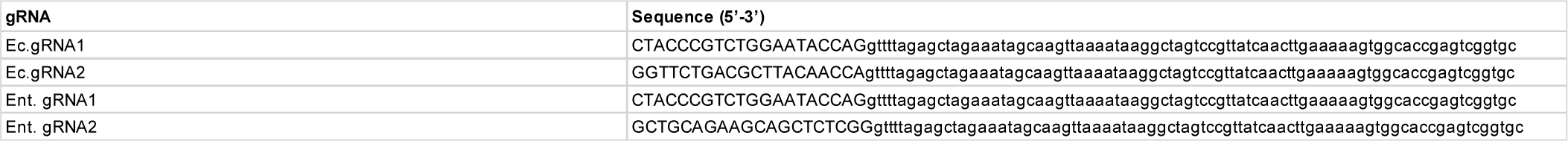
gRNA sequences tested in this study. Capital letters indicate the protospacer sequence.

**S2. Table.**
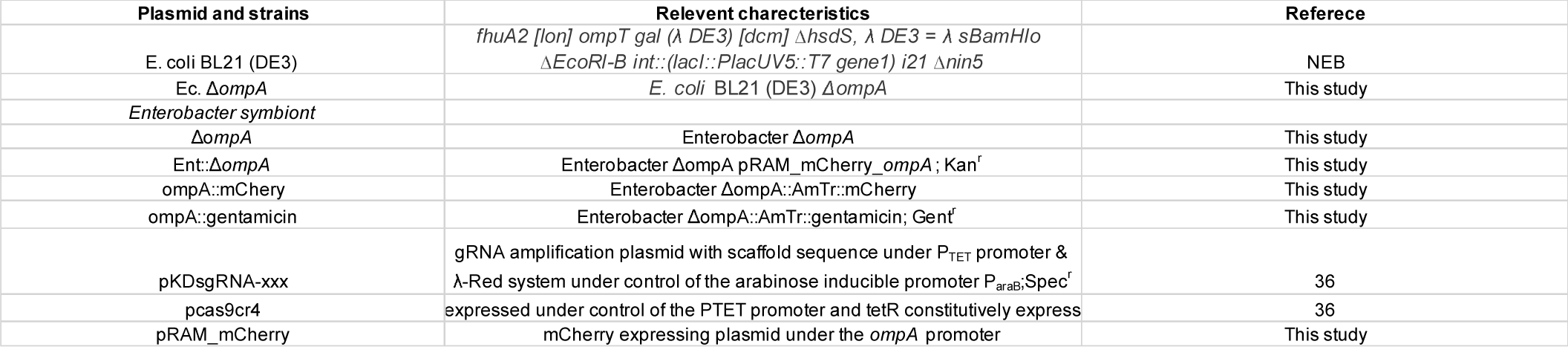
Plasmids and bacterial strains used in this study.

**S3. Table.**
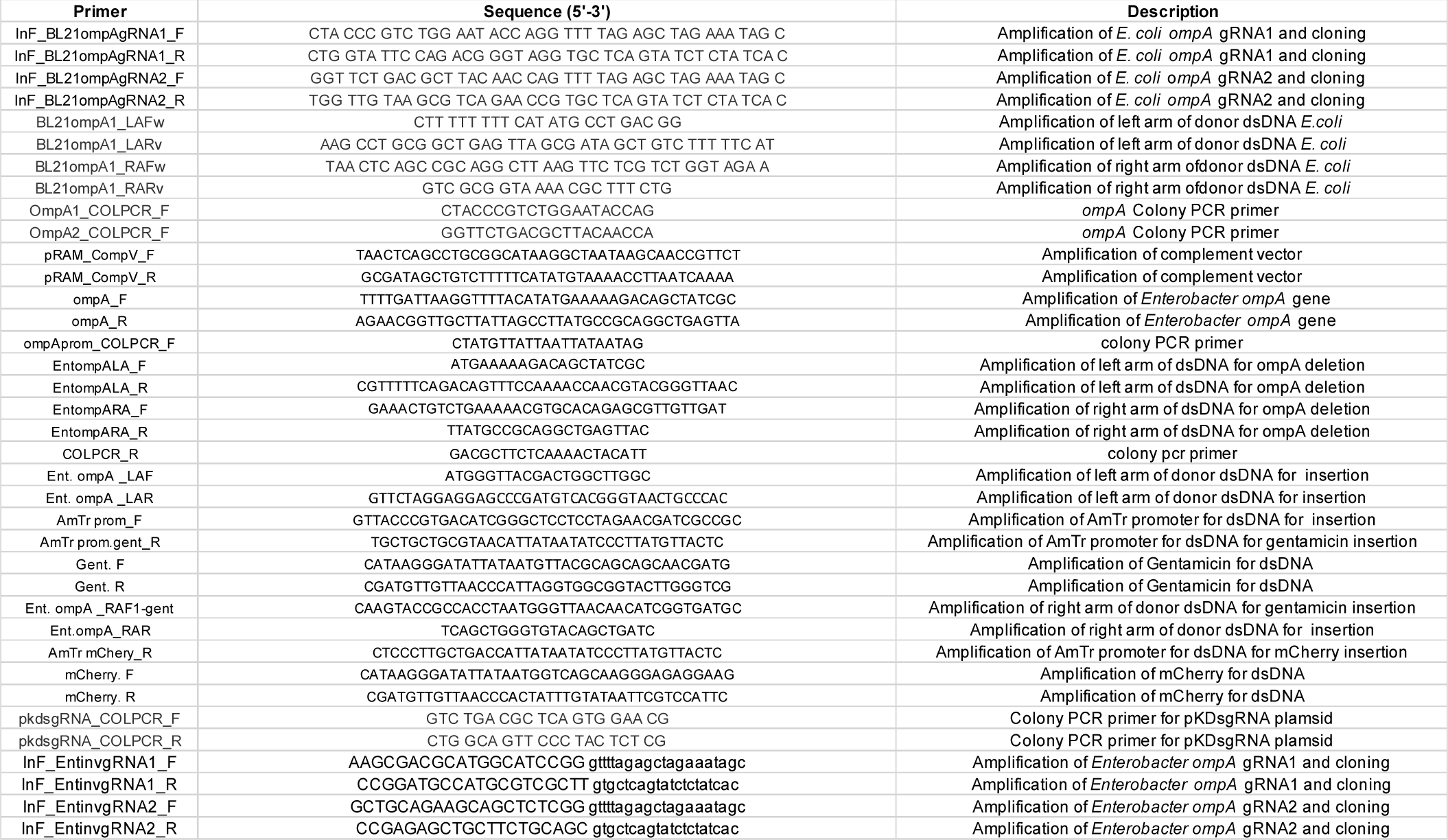
Primers used in this study.

## S1. Appendix

Multiple sequence alignment of WT and mutant *ompA* sequences of *Enterobacter* and *E. coli*.

Multiple sequence alignment *E. coli* WT *ompA* and mutant *ompA*

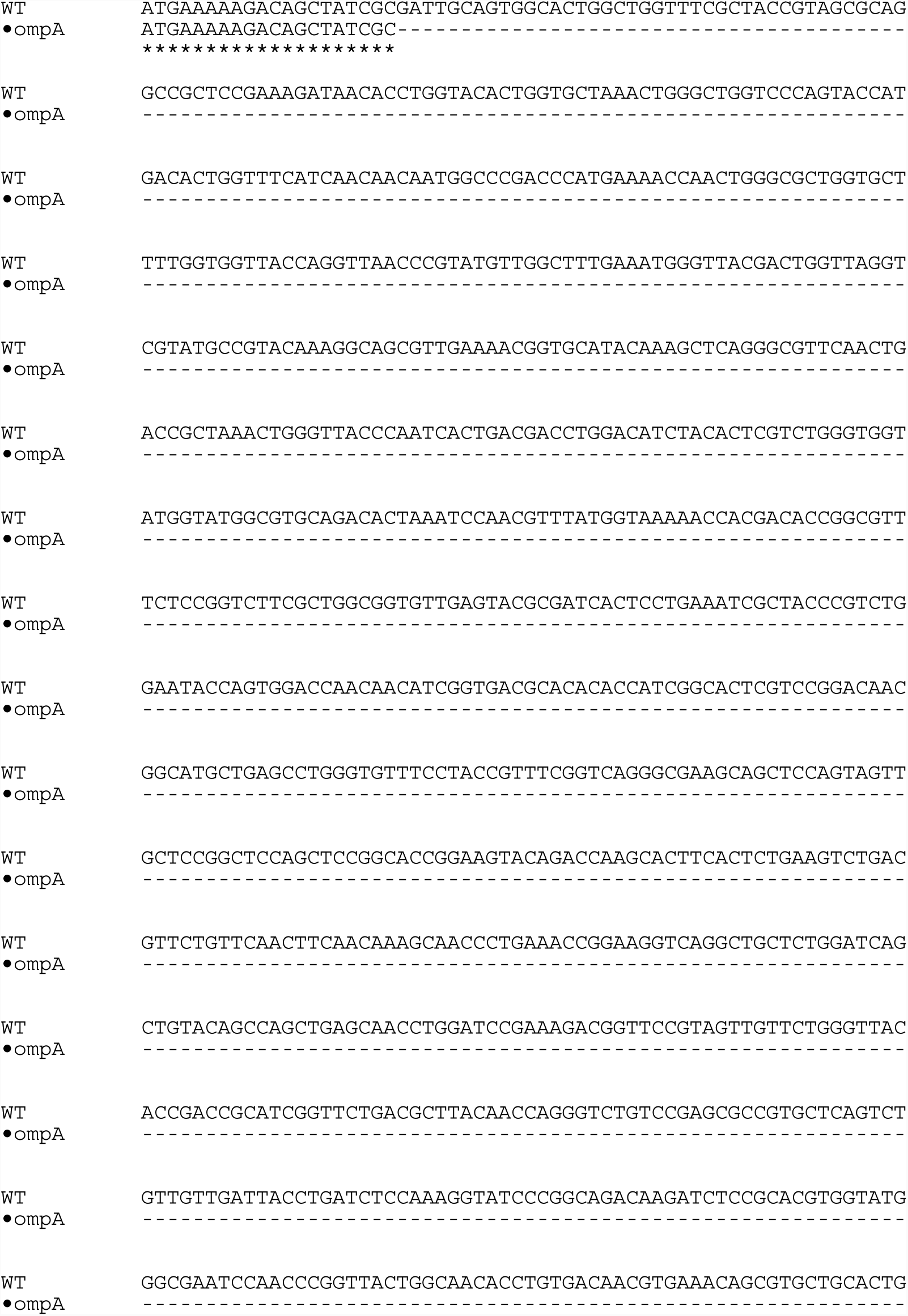
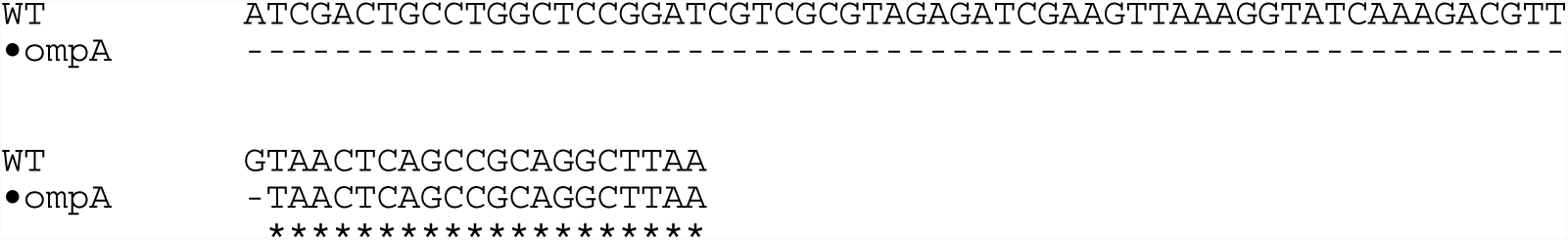

Alignment of *Enterobacter* WT *ompA* and mutant *ompA* sequence

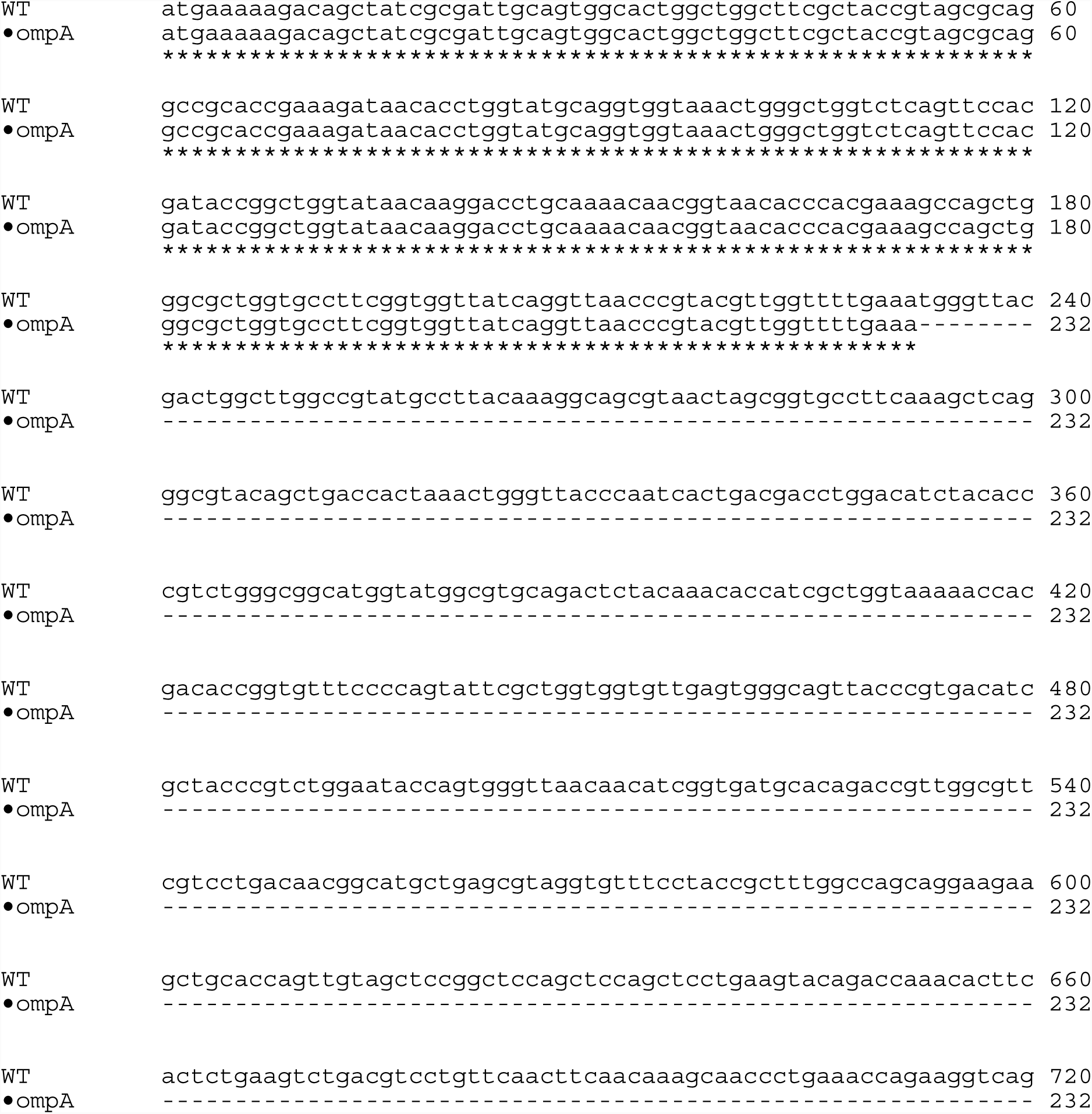
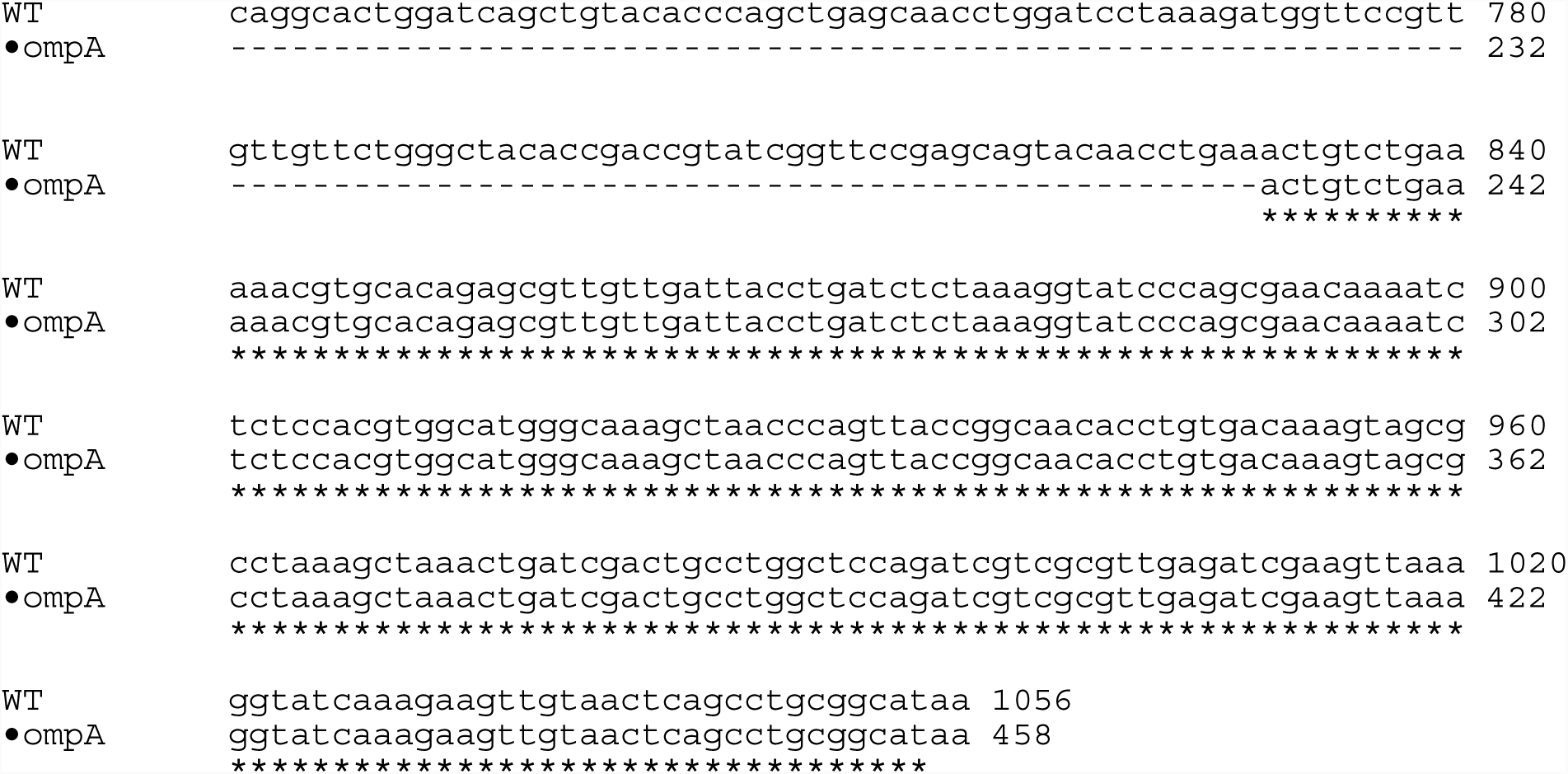

## References

1. Osei-Poku J, Mbogo CM, Palmer WJ, Jiggins FM. Deep sequencing reveals extensive variation in the gut microbiota of wild mosquitoes from Kenya. Molecular Ecology. 2012;21(20):5138–50. doi:10.1111/j.1365-294X.2012.05759.x.

2. Boissière A, Tchioffo MT, Bachar D, Abate L, Marie A, Nsango SE, et al. Midgut microbiota of the malaria mosquito vector Anopheles gambiae and interactions with Plasmodium falciparum infection. PLoS Pathogens. 2012;8(5):e1002742. doi:10.1371/journal.ppat.1002742.

3. David MR, Santos LMBD, Vicente ACP, Maciel-de-Freitas R. Effects of environment, dietary regime and ageing on the dengue vector microbiota: evidence of a core microbiota throughout Aedes aegypti lifespan. Memórias do Instituto Oswaldo Cruz. 2016;111(9):577–87. doi:10.1590/0074-02760160238.

4. Muturi EJ, Kim C-H, Bara J, Bach EM, Siddappaji MH. Culex pipiens and Culex restuans mosquitoes harbor distinct microbiota dominated by few bacterial taxa. Parasites & Vectors. 2016;9(1):18. doi:10.1186/s13071-016-1299-6.

5. Hughes GL, Dodson BL, Johnson RM, Murdock CC, Tsujimoto H, Suzuki Y, et al. Native microbiome impedes vertical transmission of Wolbachia in Anopheles mosquitoes. Proceedings of the National Academy of Sciences of the United States of America. 2014;111(34):12498–503. doi:10.1073/pnas.1408888111.

6. Coon KL, Brown MR, Strand MR. Mosquitoes host communities of bacteria that are essential for development but vary greatly between local habitats. Molecular Ecology. 2016;25(22):5806–26. doi:10.1111/mec.13877.

7. Hegde S, Khanipov K, Albayrak L, Golovko G, Pimenova M, Saldaña MA, et al. Microbiome interaction networks and community structure from laboratory-reared and field-collected Aedes aegypti, Aedes albopictus, and Culex quinquefasciatus mosquito vectors. Frontiers in Microbiology. 2018;9:715. doi:10.3389/fmicb.2018.02160.

8. Wang Y, Gilbreath TM, Kukutla P, Yan G, Xu J. Dynamic gut microbiome across life history of the malaria mosquito Anopheles gambiae in Kenya. PLoS ONE. 2011;6(9):e24767. doi:10.1371/journal.pone.0024767.

9. Weiss BL, Aksoy S. Microbiome influences on insect host vector competence. Trends in Parasitology. 2011;27(11):514–22. doi:10.1016/j.pt.2011.05.001.

10. Saldaña MA, Hegde S, Hughes GL. Microbial control of arthropod-borne disease. Memórias do Instituto Oswaldo Cruz. 2017;112(2):81–93. doi:10.1590/0074-02760160373.

11. Hegde S, Rasgon JL, Hughes GL. The microbiome modulates arbovirus transmissionin mosquitoes. Current Opinion in Virology. 2015;15:97–102. doi:10.1016/j.coviro.2015.08.011.

12. Audsley MD, Ye YH, McGraw EA. The microbiome composition of Aedes aegypti is not critical for Wolbachia-mediated inhibition of dengue virus. PLoS Neglected Tropical Diseases. 2017;11(3):e0005426. doi:10.1371/journal.pntd.0005426.

13. Muturi EJ, Bara JJ, Rooney AP, Hansen AK. Midgut fungal and bacterial microbiota of Aedes triseriatus and Aedes japonicus shift in response to La∼Crosse virus infection. Molecular Ecology. 2016;25(16):4075–90. doi:10.1111/mec.13741.

14. Kumar S, Molina-Cruz A, Gupta L, Rodrigues J, Barillas-Mury C. A peroxidase/dual oxidase system modulates midgut epithelial immunity in Anopheles gambiae. Science. 2010;327(5973):1644–8. doi:10.1126/science.1184008.

15. Pang X, Xiao X, Liu Y, Zhang R, Liu J, Liu Q, et al. Mosquito C-type lectins maintain gut microbiome homeostasis. Nature Microbiology. 2016;1(5):16023. doi:10.1038/nmicrobiol.2016.23.

16. Short SM, Mongodin EF, MacLeod HJ, Talyuli OAC, Dimopoulos G. Amino acid metabolic signaling influences Aedes aegypti midgut microbiome variability. PLoS Neglected Tropical Diseases. 2017;11(7):e0005677–29. doi:10.1371/journal.pntd.0005677.

17. Stathopoulos S, Neafsey DE, Lawniczak MKN, Muskavitch MAT, Christophides GK. Genetic dissection of Anopheles gambiae gut epithelial responses to Serratia marcescens. PLOS Pathogens. 2014;10(3):e1003897. doi:10.1371/journal.ppat.1003897.s015.

18. Xiao X, Yang L, Pang X, Zhang R, Zhu Y, Wang P, et al. A Mesh-Duox pathway regulates homeostasis in the insect gut. Nature Microbiology. 2017;2(5):17020. doi:10.1038/nmicrobiol.2017.20.

19. Powell JE, Leonard SP, Kwong WK, Engel P, Moran NA. Genome-wide screen identifies host colonization determinants in a bacterial gut symbiont. Proceedings of the National Academy of Sciences of the United States of America. 2016:201610856. doi:10.1073/pnas.1610856113.

20. Kim JK, Kwon JY, Kim S-K, Han SH, Won YJ, Lee J-H, et al. Purine biosynthesis, biofilm formation, and persistence of an snsect-microbe gut symbiosis. Appl Environ Microbiol. 2014;80(14):4374–82. doi:10.1128/AEM.00739-14.

21. Mulcahy H, Sibley CD, Surette MG, Lewenza S. Drosophila melanogaster as an animal model for the study of Pseudomonas aeruginosa biofilm infections in vivo. PLOS Pathogens. 2011;7(10):e1002299–14. doi:10.1371/journal.ppat.1002299.

22. Maltz M, Graf J. The type II secretion system is essential for erythrocyte lysis and gut colonization by the leech digestive tract symbiont Aeromonas veronii. Applied and Environmental Microbiology. 2011;77(2):597–603. doi:10.1128/AEM.01621-10.

23. Weiss BL, Wu Y, Schwank JJ, Tolwinski NS, Aksoy S. An insect symbiosis is influenced by bacterium-specific polymorphisms in outer-membrane protein A. Proceedings of the National Academy of Sciences of the United States of America. 2008;105(39):15088–93. doi:10.1073/pnas.0805666105.

24. Maltz MA, Weiss BL, Oapos Neill M, Wu Y, Aksoy S. OmpA-mediated biofilm formation is essential for the commensal bacterium Sodalis glossinidius to colonize the tsetse fly gut. Applied and Environmental Microbiology. 2012;78(21):7760–8. doi:10.1128/AEM.01858-12.

25. Riehle MA, Jacobs-Lorena M. Using bacteria to express and display anti-parasite molecules in mosquitoes: current and future strategies. Insect Biochemistry and Molecular Biology. 2005;35(7):699–707. doi:10.1016/j.ibmb.2005.02.008.

26. Dennison NJ, Saraiva RG, Cirimotich CM, Mlambo G, Mongodin EF, Dimopoulos G. Functional genomic analyses of Enterobacter, Anopheles and Plasmodium reciprocal interactions that impact vector competence. Malaria Journal. 2016;15(1):425. Epub 3. doi:10.1186/s12936-016-1468-2.

27. Pei D, Jiang J, Yu W, Kukutla P, Uentillie A, Xu J. The waaL gene mutation compromised the inhabitation of Enterobacter sp. Ag1 in the mosquito gut environment. Parasites & Vectors. 2015:1–10. doi:10.1186/s13071-015-1049-1.

28. Enomoto S, Chari A, Clayton AL, Dale C. Quorum sensing attenuates virulence in Sodalis praecaptivus. Cell Host & Microbe. 2017;21(5):629–36.e5. doi:10.1016/j.chom.2017.04.003.

29. Dale C, Young SA, Haydon DT, Welburn SC. The insect endosymbiont Sodalis glossinidius utilizes a type III secretion system for cell invasion. Proceedings of the National Academy of Sciences. 2001;98(4):1883–8. doi:10.1073/pnas.021450998.

30. Selle K, microbiology RBTi, 2015. Harnessing CRISPR–Cas systems for bacterial genome editing. Elsevier. doi:10.1016/j.tim.2015.01.008.

31. Sander JD, Joung JK. CRISPR-Cas systems for editing, regulating and targeting genomes. Nature Biotechnology. 2014;32(4):347–55. doi:10.1038/nbt.2842.

32. Barrangou R, van Pijkeren JP. Exploiting CRISPR-Cas immune systems for genome editing in bacteria. Current Opinion in Biotechnology. 2016;37:61–8. doi:10.1016/j.copbio.2015.10.003.

33. Jiang Y, Chen B, Duan C, Sun B, Yang J, Yang S. Multigene editing in the Escherichia coli genome via the CRISPR-Cas9 system. Applied and Environmental Microbiology. 2015;81(7):2506–14. doi:10.1128/AEM.04023-14.

34. Jiang W, Bikard D, Cox D, Zhang F, Marraffini LA. RNA-guided editing of bacterial genomes using CRISPR-Cas systems. Nature Biotechnology. 2013;31(3):233–9. doi:10.1038/nbt.2508.

35. Reisch CR, Prather KLJ. The no-SCAR (Scarless Cas9 Assisted Recombineering) system for genome editing in Escherichia coli. Scientific Reports. 2015;5:15096. doi:10.1038/srep15096.

36. Li Y, Lin Z, Huang C, Zhang Y, Wang Z, Tang Y-j, et al. Metabolic engineering of Escherichia coli using CRISPR–Cas9 meditated genome editing. Metabolic Engineering. 2015;31:13–21. doi:10.1016/j.ymben.2015.06.006.

37. Ronda C, Pedersen LE, Sommer MOA, Nielsen AT. CRMAGE: CRISPR Optimized MAGE Recombineering. Scientific Reports. 2016;6(1):1200. doi:10.1038/srep19452.

38. Tong Y, Robertsen HL, Blin K, Weber T, Lee SY. CRISPR-Cas9 Toolkit for Actinomycete Genome Editing. Methods in molecular biology 2018;1671(1):163–84. doi:10.1007/978-1-4939-7295-1_11.

39. Oh J-H, van Pijkeren JP. CRISPR-Cas9-assisted recombineering in Lactobacillus reuteri. Nucleic Acids Research. 2014;42(17):e131–e. doi:10.1093/nar/gku623.

40. Mougiakos I, Bosma EF, Weenink K, Vossen E, Goijvaerts K, van der Oost J, et al. Efficient genome editing of a facultative thermophile using mesophilic spCas9. ACS Synthetic Biology. 2017;6(5):849–61. doi:10.1021/acssynbio.6b00339.

41. Li K, Cai D, Wang Z, He Z, Chen S. Development of an efficient genome editing tool in Bacillus licheniformis ssing CRISPR-Cas9 nickase. Applied and Environmental Microbiology. 2018:AEM.02608–17. doi:10.1128/AEM.02608-17.

42. Jiang Y, Qian F, Yang J, Liu Y, Dong F, Xu C, et al. CRISPR-Cpf1 assisted genome editing of Corynebacterium glutamicum. Nat Commun. 2017;8:15179. Epub 2017/05/05. doi:10.1038/ncomms15179.

43. Cobb RE, Wang Y, Zhao H. High-efficiency multiplex genome editing of Streptomyces species using an engineered CRISPR/Cas system. ACS Synth Biol. 2015;4(6):723–8. Epub 2014/12/03. doi:10.1021/sb500351f.

44. Waller MC, Bober JR, Nair NU, Beisel CL. Toward a genetic tool development pipeline for host-associated bacteria. Current Opinion in Microbiology. 2017;38:156–64. doi:10.1016/j.mib.2017.05.006.

45. Wilke ABB, Marrelli MT. Paratransgenesis: a promising new strategy for mosquito vector control. Parasites & Vectors. 2015;8(1):391–19. doi:10.1186/s13071-015-0959-2.

46. Arora AK, Douglas AE. Hype or opportunity? Using microbial symbionts in novel strategies for insect pest control. Journal of insect physiology. 2017;103:10–7. doi:10.1016/j.jinsphys.2017.09.011.

47. Ricci I, Damiani C, Capone A, DeFreece C, Rossi P, Favia G. Mosquito/microbiota interactions: from complex relationships to biotechnological perspectives. Current Opinion in Microbiology. 2012;15(3):278–84. doi:10.1016/j.mib.2012.03.004.

48. Beard CB, Mason PW, Aksoy S, Tesh RB, Richards FF. Transformation of an insect symbiont and expression of a foreign gene in the Chagas’ disease vector Rhodnius prolixus. American Journal of Tropical Medicine and Hygiene. 1992;46(2):195–200.

49. Bisi DC, Lampe DJ. Secretion of anti-Plasmodium effector proteins from a natural Pantoea agglomerans isolate by using PelB and HlyA secretion signals. Applied and Environmental Microbiology. 2011;77(13):4669–75. doi:10.1128/AEM.00514-11.

50. Hughes GL, Allsopp PG, Webb RI, Yamada R, Iturbe-Ormaetxe I, Brumbley SM, et al. Identification of yeast associated with the planthopper, Perkinsiella saccharicida: potential applications for Fiji leaf gall control. Current Microbiology. 2011;63(4):392–401. doi:10.1007/s00284-011-9990-5.

51. Medina F, Li H, Vinson SB, Coates CJ. Genetic transformation of midgut bacteria from the red imported fire ant (Solenopsis invicta). Current Microbiology. 2009;58(5):478–82. doi:10.1007/s00284-008-9350-2.

52. Bextine B, Lauzon C, Potter S, Lampe D, Miller TA. Delivery of a genetically marked Alcaligenes sp. to the glassy-winged sharpshooter for use in a paratransgenic control strategy. Current Microbiology. 2004;48(5):327–31. doi:10.1007/s00284-003-4178-2.

53. Hurwitz I, Hillesland H, Fieck A, Das P, Durvasula R. The paratransgenic sand fly: A platform for control of Leishmania transmission. Parasites & Vectors. 2011;4(1):82. doi:10.1186/1756-3305-4-82.

54. Durvasula RV, Gumbs A, Panackal A, Kruglov O, Taneja J, Kang AS, et al. Expression of a functional antibody fragment in the gut of Rhodnius prolixus via transgenic bacterial symbiont Rhodococcus rhodnii. Medical and Veterinary Entomology. 1999;13(2):115–9.

55. Wang S, Ghosh AK, Bongio N, Stebbings KA, Lampe DJ, Jacobs-Lorena M. Fighting malaria with engineered symbiotic bacteria from vector mosquitoes. Proceedings of the National Academy of Sciences of the United States of America. 2012;109(31):12734–9. doi:10.1073/pnas.1204158109.

56. Wu SC-Y, Maragathavally KJ, Coates CJ, Kaminski JM. Steps toward targeted insertional mutagenesis with class II transposable elements. Methods in molecular biology 2008;435:139–51. doi:10.1007/978-1-59745-232-8_10.

57. Tikhe CV, Martin TM, Howells A, Delatte J, Husseneder C. Assessment of genetically engineered Trabulsiella odontotermitis as a ‘Trojan Horse’ for paratransgenesis in termites. BMC Microbiology. 2016;16(1):355. doi:10.1186/s12866-016-0822-4.

58. Pittman GW, Brumbley SM, Allsopp PG, Oapos Neill SL. Assessment of gut bacteria for a paratransgenic approach to control Dermolepida albohirtum larvae. Applied and Environmental Microbiology. 2008;74(13):4036–43. doi:10.1128/AEM.02609-07.

59. Pontes MH, Dale C. Lambda red-mediated genetic modification of the insect endosymbiont Sodalis glossinidius. Applied and Environmental Microbiology. 2011;77(5):1918–20. doi:10.1128/AEM.02166-10.

60. Wang S, Dos-Santos ALA, Huang W, Liu KC, Oshaghi MA, Wei G, et al. Driving mosquito refractoriness to Plasmodium falciparum with engineered symbiotic bacteria. Science (New York, NY). 2017;357(6358):1399–402. doi:10.1126/science.aan5478.

61. Dotson EM, Plikaytis B, Shinnick TM, Durvasula RV, Beard CB. Transformation of Rhodococcus rhodnii, a symbiont of the Chagas disease vector Rhodnius prolixus, with integrative elements of the L1 mycobacteriophage. Infection, genetics and evolution. 2003;3(2):103–9.

62. Bextine B, Lampe D, Lauzon C, Jackson B, Miller TA. Establishment of a genetically marked insect-derived symbiont in multiple host plants. Curr Microbiol. 2005;50(1):1–7. doi:10.1007/s00284-004-4390-8.

63. Wu P, Sun P, Nie K, Zhu Y, Shi M, Xiao C, et al. A gut commensal bacterium promotes mosquito permissiveness to arboviruses. Cell Host & Microbe. 2019;25(1):101–12.e5. doi:10.1016/j.chom.2018.11.004.

64. Coon KL, Vogel KJ, Brown MR, Strand MR. Mosquitoes rely on their gut microbiota for development. Molecular Ecology. 2014;23(11):2727–39. doi:10.1111/mec.12771.

65. Lindh JM, Borg-Karlson AK, Faye I. Transstadial and horizontal transfer of bacteria within a colony of Anopheles gambiae (Diptera: Culicidae) and oviposition response to bacteria-containing water. Acta Tropica. 2008;107(3):242–50. doi:10.1016/j.actatropica.2008.06.008.

66. Coon KL, Brown MR, Strand MR. Gut bacteria differentially affect egg production in the anautogenous mosquito Aedes aegypti and facultatively autogenous mosquito Aedes atropalpus (Diptera: Culicidae). Parasites & Vectors. 2016;9(1):375. doi:10.1186/s13071-016-1660-9.

67. Ribet D, Cossart P. How bacterial pathogens colonize their hosts and invade deeper tissues. Microbes and infection. 2015;17(3):173–83. doi:10.1016/j.micinf.2015.01.004.

68. Ramirez JL, Short SM, Bahia AC, Saraiva RG, Dong Y, Kang S, et al. Chromobacterium Csp_P reduces malaria and dengue infection in vector mosquitoes and has entomopathogenic and in vitro anti-pathogen activities. PLoS Pathogens. 2014;10(10):e1004398. doi:10.1371/journal.ppat.1004398.

69. Valzania L, Martinson VG, Harrison RE, Boyd BM, Coon KL, Brown MR, et al. Both living bacteria and eukaryotes in the mosquito gut promote growth of larvae. PLoS Neglected Tropical Diseases. 2018;12(7):e0006638. doi:10.1371/journal.pntd.0006638.

70. Correa MA, Matusovsky B, Brackney DE, Steven B. Generation of axenic Aedes aegypti demonstrate live bacteria are not required for mosquito development. Nature Communications. 2018;9(1):R37. doi:10.1038/s41467-018-07014-2.

71. Favia G, Ricci I, Damiani C, Raddadi N, Crotti E, Marzorati M, et al. Bacteria of the genus Asaia stably associate with Anopheles stephensi, an Asian malarial mosquito vector. Proceedings of the National Academy of Sciences. 2007;104(21):9047–51. doi:10.1073/pnas.0610451104.

72. Chaverra-Rodriguez D, Macias VM, Hughes GL, Pujhari S, Suzuki Y, Peterson DR, et al. Targeted delivery of CRISPR-Cas9 ribonucleoprotein into arthropod ovaries for heritable germline gene editing. Nature Communications. 2018;9(1):245. doi:10.1038/s41467-018-05425-9.

73. Li M, Bui M, Yang T, Bowman CS, White BJ, Akbari OS. Germline Cas9 expression yields highly efficient genome engineering in a major worldwide disease vector,Aedes aegypti. Proceedings of the National Academy of Sciences of the United States of America. 2017;114(49):E10540–E9. doi:10.1073/pnas.1711538114.

74. Montague TG, Cruz JM, Gagnon JA, Church GM, Valen E. CHOPCHOP: a CRISPR/Cas9 and TALEN web tool for genome editing. Nucleic Acids Research. 2014;42(W1):W401–W7. doi:10.1093/nar/gku410.

75. Labun K, Montague TG, Gagnon JA, Thyme SB, Valen E. CHOPCHOP v2: a web tool for the next generation of CRISPR genome engineering. Nucleic Acids Research. 2016;44(W1):W272–W6. doi:10.1093/nar/gkw398.

76. Trehan A, Kiełbus M, Czapinski J, Stepulak A, Huhtaniemi I, Rivero-Müller A. REPLACR-mutagenesis, a one-step method for site-directed mutagenesis by recombineering. Scientific Reports. 2015:1–9. doi:10.1038/srep19121.

77. Kozlova EV, Khajanchi BK, Popov VL, Wen J, Chopra AK. Impact of QseBC system in c-di-GMP-dependent quorum sensing regulatory network in a clinical isolate SSU of Aeromonas hydrophila. Microbial pathogenesis. 2012;53(3-4):115–24. doi:10.1016/j.micpath.2012.05.008.

78. O’Toole GA, Kolter R. Flagellar and twitching motility are necessary for Pseudomonas aeruginosa biofilm development. Molecular Microbiology. 1998;30(2):295–304. doi:10.1046/j.1365-2958.1998.01062.x.

