## Supplementary figures and images for "CRISPR/Cas9-mediated gene deletion of the *ompA gene* in symbiotic *Enterobacter* impairs biofilm formation and reduces gut colonization of *Aedes aegypti* mosquitoes"

### Figure S1

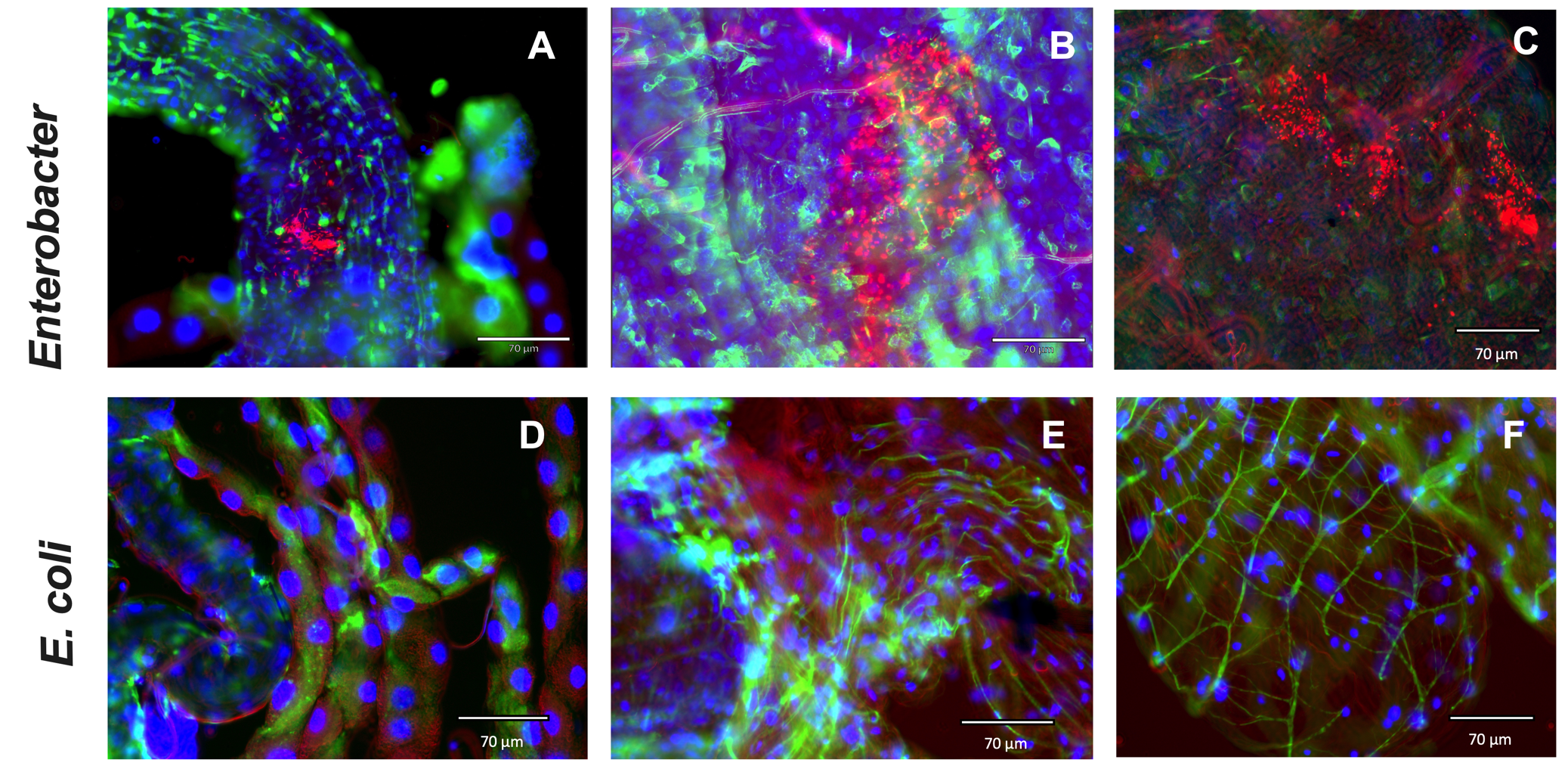
